# Organization of Upstream ESCRT Machinery at the HIV-1 Budding Site

**DOI:** 10.1101/2022.09.13.507863

**Authors:** Arpa Hudait, James H. Hurley, Gregory A. Voth

## Abstract

In the late stages of the HIV-1 life cycle, membrane localization and self-assembly of the Gag polyproteins induce membrane deformation and budding. However, release of the immature virion requires direct interaction between Gag lattice and upstream ESCRT machinery at the budding site, followed by assembly of the downstream ESCRT-III factors, culminating in membrane scission. In this work, using “bottom-up” coarse-grained (CG) molecular dynamics (MD) simulations we investigated the interactions between Gag and different upstream ESCRT components to delineate the molecular organization of proteins at the membrane neck of the HIV-1 budding site. We developed CG models of upstream ESCRT proteins and HIV-1 structural protein Gag based on experimental structural data and extensive all-atom MD simulations. We find that ESCRT-I proteins bound to the immature Gag lattice can recruit multiple copies of ESCRT-II coating the membrane neck. ESCRT-I can effectively oligomerize to higher-order complexes both in absence of ESCRT-II and when multiple copies of ESCRT-II are localized at the bud neck. The ESCRT-I/II supercomplexes observed in our simulations exhibit predominantly extended conformations. Importantly, the ESCRT-I/II supercomplex modulates the membrane mechanical properties at the budding site by decreasing the overall Gaussian curvature of membrane neck. Our findings serve to elucidate a network of interactions between the upstream ESCRT machinery, immature Gag lattice, and membrane bud neck that regulate the protein assemblies and enable bud neck constriction.

## Introduction

A critical step in the HIV-1 life cycle is the assembly of viral Gag polyproteins at the cellular membrane, followed by a series of membrane remodeling events at the site of viral protein localization that culminates in the budding and fission of the immature virion (1-4). To aid in the viral particle release, HIV-1 recruits the endosomal sorting complexes required for transport (ESCRT) machinery (5-8). Multiple studies have demonstrated significant inhibition of viral particle production when immature virions are unable to recruit the ESCRT machinery (9-12). The ESCRT machinery facilitates membrane scission in a variety of other cellular processes such as the formation of multivesicular bodies (13), nuclear envelope reformation (14,15), and cytokinetic membrane scission (16,17). ESCRTs predominantly direct the budding and scission of the membrane neck pointing away (reverse-topology) from the cytoplasm. Recent studies have shown that the ESCRT complexes can also facilitate regular-topology scission (18-20), thus demonstrating versatility in their mechanism of action.

The initial events of immature virion formation involve steady recruitment of the HIV-1 structural protein Gag at the plasma membrane (5,21,22). The matrix (MA) domain of Gag polyprotein mediates interactions with the plasma membrane (23,24). The capsid (CA) and spacer peptide 1 (SP1) domains of the Gag provide key protein-protein contacts for assembly of the hexameric bundle, the building block of immature Gag lattice (25-27). Nucleation and growth of the Gag lattice result in a quasi-spherical shaped immature virion (28-30). Recruitment of the ESCRT proteins at the viral assembly site closely follows the Gag accumulation and oligomerization events (5,7,22,31,32). Gag polyproteins recruit the ESCRT machinery through motifs in the C-terminal p6 domain that points inwards to the center of the spherical shell. Specifically, the PTAP motif interacts with the ubiquitin E2 variant (UEV) domain of the ESCRT-I complex (10-12,33-35) and the LYPX_n_L motif interacts with the ALG-2 interacting protein X (ALIX) (36-40). When bound to the Gag lattice, headpiece region of ESCRT-I oligomerizes to a 12-membered ring facilitated by electrostatic interactions from the residues in the VPS28 N-terminal domain (NTD) (41). We have previously shown in that work using coarse-grained (CG) molecular dynamics (MD) simulations that ESCRT-I oligomerization is facilitated by optimal geometry of the immature Gag lattice. In this “geometry-dependent checkpoint” model higher-order ESCRT-I oligomers are formed at the late stages of Gag assembly when the dimension of the lumen of the growing Gag lattice approaches ∼50 nm, the outer diameter of the ESCRT-I ring (41).

ESCRT-I binds to the ESCRT-II, a peripheral membrane targeting complex through the VPS28 C-terminal domain (CTD) in the headpiece region (42,43). Next, the VPS25 subunits of the ESCRT-II then recruit the CHMP6, the most upstream component of the ESCRT-III family, also a membrane targeting protein (44,45). Overall, ESCRT-I, ESCRT-II, and ALIX constitute the upstream ESCRT machinery. ESCRT-I/II with CHMP6 provides one pathway for ESCRT-III nucleation and polymerization, while ALIX offers a secondary pathway by directly binding to the ESCRT-III components leading to virion budding (32,46-49). It is established that upstream ESCRTs play an essential role in regulating the downstream ESCRT-III self-assembly.

Membrane-protein interactions and membrane remodeling events are central to the immature virion assembly, budding and release. HIV-1 structural protein Gag binds to the membrane through a PIP2 targeting highly basic region (HBR) and is anchored to the membrane leaflet through insertion of the N-terminal myristoyl moiety (24,50). Early stages of Gag localization and hexamer nucleation can generate PIP2 enriched nanodomains in the vicinity and spontaneous membrane curvature, which in turn can localize more Gag proteins and facilitate immature lattice growth (51-53). Upstream ESCRT-I, ESCRT-II, and CHMP6 are recruited to the budding site as the immature Gag lattice grows to a spherical shell (32). ESCRT-II is a peripheral membrane protein complex consisting of two membrane targeting sites: a highly basic *α*-helix motif and a phosphatidylinositol lipid targeting GLUE domain (42). CHMP6 association with the membrane is mediated by the N-terminal basic region and myristoyl anchoring (45). Further downstream, ESCRT-III associates with the membrane and self-assembles to filaments adopting multiple shapes such as spirals, helical tubes, and cones (54-60). Theoretical studies have proposed that membrane neck scission is caused by constricting forces exerted by shape transition of ESCRT-III polymers coating the bud neck (61,62). At the bud neck, upstream ESCRT machinery bridges the immature Gag lattice and downstream ESCRT-III components. How upstream ESCRTs sense local curvature to localize at the bud neck, and influence bud neck curvature, remains an open question. Previous experimental and theoretical studies had reported that upstream ESCRTs can induce membrane deformation by localizing at the membrane neck in large concentrations facilitating vesicle formation (13,63,64). Other studies using analytical theory and continuum models demonstrated that upstream ESCRT complexes can generate spontaneous membrane curvature and budding (65,66).

In this work, we utilize CG MD simulations to investigate the higher-order organization of upstream ESCRT-I and ESCRT-II at the HIV-1 budding site and develop a composite model of the Gag and upstream ESCRT assemblies at the viral budding site. First, we derived bottom-up implicit-solvent CG models of ESCRT-II in addition to the ESCRT-I and Gag CG models developed previously (41), allowing us to model the full upstream ESCRT machinery with the immature virion. Here, the term “bottom-up” means that the CG models are systematically derived from the underlying atomistic-level interactions, as opposed to developed in an *ad hoc*, “top-down” fashion (67). We investigated ESCRT-II recruitment dynamics for two scenarios when in the initial configuration the ESCRT-I proteins bound to the immature Gag lattice are fully assembled to a 12-membered ring and instead when ESCRT-I is fully disassembled. We find that multiple ESCRT-II proteins bind to the ESCRT-I coating the bud neck. Furthermore, ESCRT-II recruitment does not significantly impact the ESCRT-I assembly behavior despite crowding at the bud neck. Our analysis also indicates that ESCRT-I/II supercomplex formation at the bud neck can modulate the bud neck Gaussian curvature and bending stiffness depending on the membrane–protein adhesion strength.

## Results

### Coarse-Grained Model of ESCRT-II Supported by Atomistic MD Simulations

ESCRT-II is a Y-shaped heterotetramer complex consisting of one VPS22 and VPS36 subunit, and two VPS25 subunits (42,68). The two VPS25 subunits each form an arm of the Y-shaped complex, while the VPS22:VPS36 subcomplex constitutes the stalk (**Fig. 1*A***). The VPS36 subunit is capped by the membrane targeting N-terminal (GLUE) domain (VPS36: residues 1-132), while the second membrane associating VPS22-H0 helix (VPS22: residues 8-23) is in the stalk. To establish the membrane-bound conformation of ESCRT-II we performed 2000 ns long all-atom (AA) MD simulations (3 replicas) of the full-length ESCRT-II binding to a PIP2 containing model membrane (**Fig. 1*A* and 1*B***). In the initial configuration, the ESCRT-II complex was positioned near the bilayer, with the lipid-binding basic patch of the GLUE domain and VPS22-H0 helix facing toward the surface of the membrane. The AA MD simulation setup is described in the *Methods*. In all the simulations, the membrane-targeting basic pockets of the GLUE domain and the basic VPS22-H0 helix make persistent PIP2 contacts (**Fig. S1** of Supporting Information). Our analysis reveals that the basic region (VPS22: residues 53-65) also forms stable contacts with the PIP2 lipids and provides an additional membrane binding interface for the stalk of the ESCRT-II complex. To characterize the orientation of the membrane-bound ESCRT-II complex we calculated the distance of the center of each domain to the membrane center from the AA MD trajectories (**Fig. 1*C***). The stalk of the ESCRT-II complex lays flat on the membrane, bound to the membrane by multiple PIP2-binding sites. Our analysis shows that the C-terminal domain of the VPS36 subunit and the VPS25 subunit contacting the C-terminal domain of the VPS36 subunit orients away from the membrane. The other VPS25 subunit contacting the C-terminal domain of the VPS22 remains closely associated with the membrane despite not targeting PIP2 lipids. Taken together, the AA MD simulations indicate that the full-length ESCRT-II complex effectively targets membrane using a multivalent mechanism, i.e., simultaneously engaging multiple PIP2-binding sites.

**Fig. 1.**
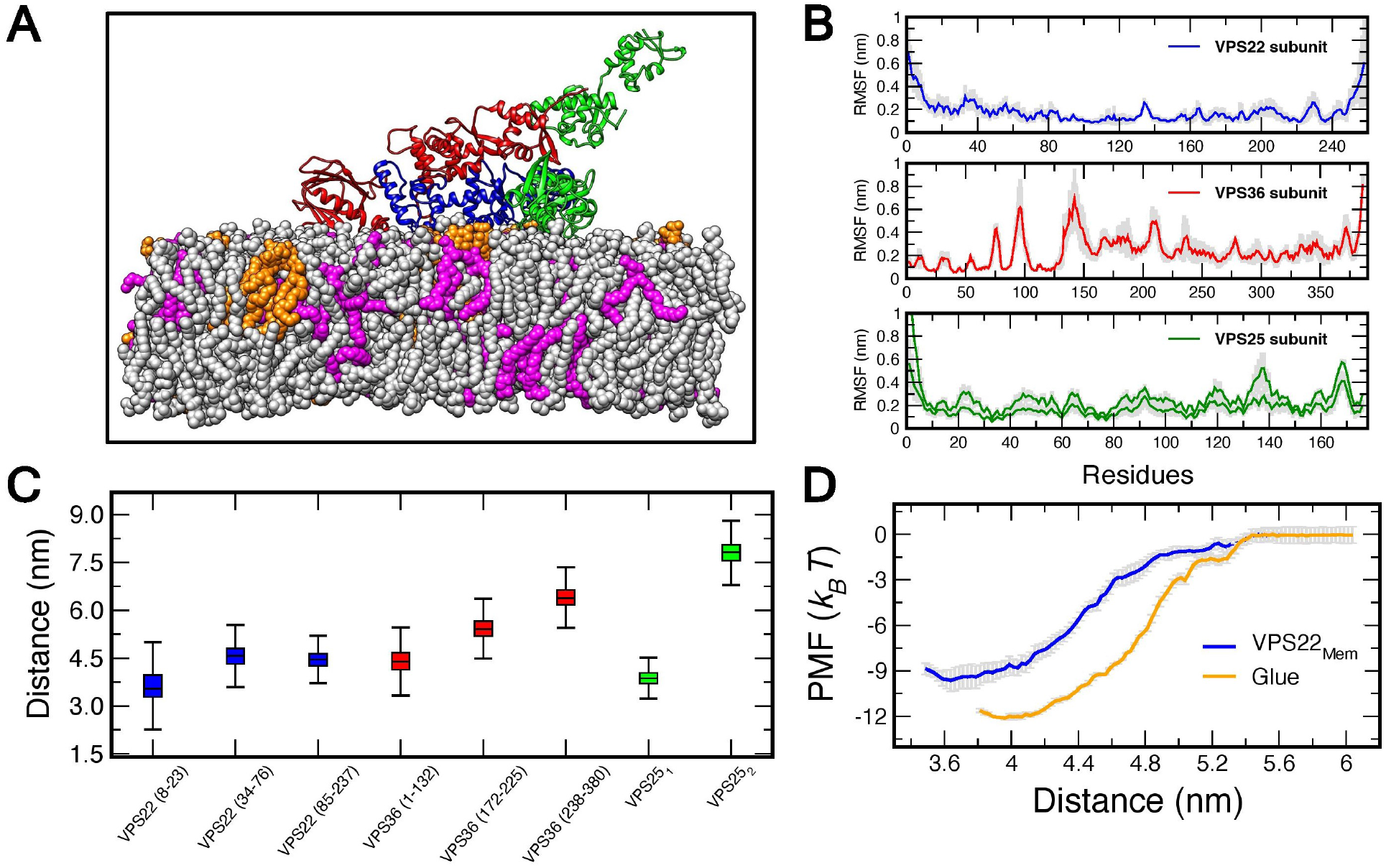
All-atom MD simulations of the ESCRT-II complex bound to the membrane. (*A*) Ribbon representation of the full-length ESCRT-II complex bound to POPC(76%):POPE (20%):PIP2(4%) membrane. The POPC, POPE and PIP2 lipids are shown in silver, magenta and orange spheres, respectively. VPS22, VPS36 and VPS25 subunits of ESCRT-II are shown in blue, red and green ribbons, respectively. (*B*) RMSFs of individual residues for each subunit of the ESCRT-II complex. The solid line represents the mean, and the shaded region represents standard deviation. The mean and standard deviation for each residue is calculated from final 1000 ns of each replica trajectory. Analysis of the RMSF reveals that the *β*6/*β*7 loop (VPS36: 87-105), VPS36 linker (VPS36: 132-171) and the C-terminal WH2 domain of VPS25 are particularly plastic. (*C*) Box plots of the distance between the center of mass of ESCRT-II residues (specified in parentheses) to the membrane center. The box plots are prepared from final 1000 ns of each replica trajectory. (*D*) Potential of mean force (PMF) for GLUE monomer and VPS22_Mem_ binding to the membrane. Error bars (shown in silver in the background) indicate standard deviation of the PMF calculated from block analysis.

We then derived a bottom-up CG model of the ESCRT-II complex from the membrane-bound conformations generated from AA MD trajectories. First, the CG model of ESCRT-II was mapped from the AA MD trajectories. The CG model of ESCRT-II complex contains 222 CG sites with an average resolution of 4.5 amino acid residues per site (**Fig. 2*A***). To capture the intraprotein flexibility, bonded topology of the CG model and the harmonic force constant of these bonds were derived from the AA MD trajectory. The details of the CG model development are provided in *Methods*. We then identified CG sites of the ESCRT-II that corresponds to amino acids that make persistent PIP2 contacts in the AA MD simulations. Attractive interactions (*E*_Esc-II/Mem_) were added between these CG sites of the ESCRT-II and the headgroup of the CG lipid to model the membrane association of the ESCRT-II complex. In the CG MD simulations the membrane was modeled with a three-site quasi-monolayer highly CG’ed lipid (69). The purpose of the latter over a more highly resolved CG lipid model is to sufficiently increase the computational efficiency of the simulation while retaining adequate physical accuracy.

**Fig. 2.**
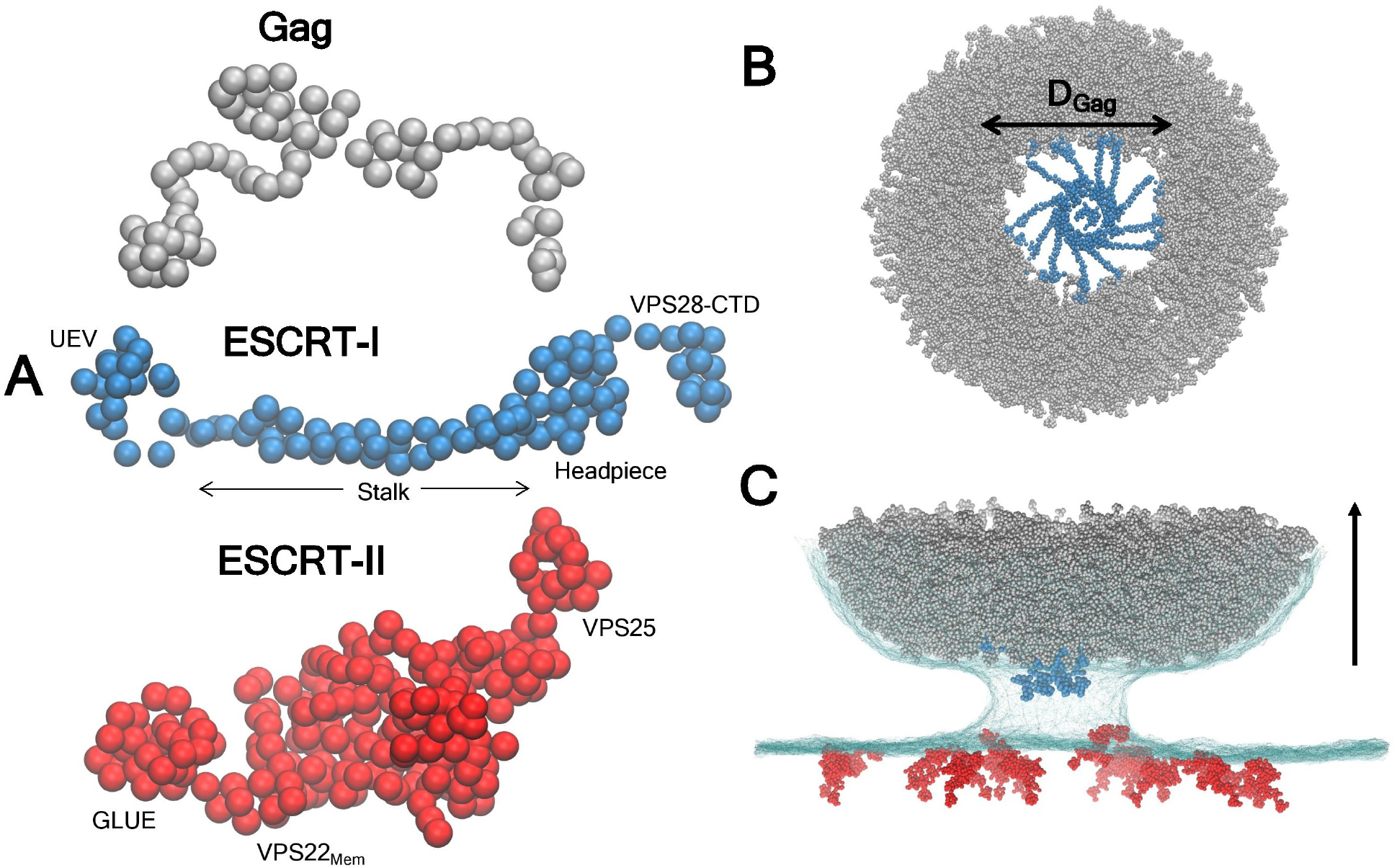
Overview of the CG models and CG MD simulation setup. (*A*) CG representation of Gag (78 sites), ESCRT-I (109 sites), and ESCRT-II (222 sites). Key regions of each protein are labeled. (*B*) Top view (looking down from the bud neck to the virion aperture) of the 12-mer ESCRT-I ring docked at the Gag shell of diameter (*D*_Gag_) 50 nm. (C) A representative snapshot of an initial CG MD simulation configuration. The Gag shell is shown in gray, ESCRT-I shown in blue, and ESCRT-II in red sphere. Initially, 16 copies of ESCRT-II are randomly distributed at the flat section of the membrane. The membrane is shown in cyan mesh. The upward arrow indicates direction of the viral budding, i.e., budding away from the cell.

To obtain estimates for the membrane association interaction strength of the CG ESCRT-II we first quantified the binding affinity of the membrane targeting domains of the all-atom ESCRT-II complex. To calculate the binding affinity, we constructed potential of mean force (PMF) profiles of protein binding and unbinding from all-atom umbrella sampling (US) simulations (details of the US simulation procedure are provided in *Methods*). For the US simulation, we used the distance between the center of lipid bilayer and protein as the reaction coordinate. We note that calculating the PMF profiles of the full-length ESCRT-II complex in AA MD simulations is prohibitively expensive due to the overall system size. Hence, for the AA US simulations we considered the GLUE domain and the membrane targeting region of the VPS22 subunit (VPS22_Mem_). Here, VPS22_Mem_ (VPS22: residues 1-76) encompasses both membrane binding regions of VPS22 as observed in the AA MD simulations of the full-length ESCRT-II complex. We approximate that the binding affinity of the full-length ESCRT-II complex is a combination of the contributions from the GLUE and VPS22_Mem_ regions. The calculated binding affinity is −12 *k*_*B*_*T* for GLUE domain and −10 *k*_*B*_*T* for VPS22_Mem_ (**Fig. 1*D***). Our results demonstrate that both membrane targeting regions of the ESCRT-II complex exhibit comparable membrane binding affinity, and agree with liposome binding assay experiments of ESCRT-II deletion constructs (42). Finally, using the AA binding affinity estimate we evaluated the appropriate *E*_Esc-II/Mem_ for the CG ESCRT-II model. To do so, we varied *E*_Esc-II/Mem_ and calculated the binding affinity of CG ESCRT-II to the membrane for each *E*_Esc-II/Mem_ from US simulations (details of the CG US simulations provided in *Methods*). We find that at *E*_Esc-II/Mem_ = 3.0 kcal/mol the binding affinity (−20 *k*_*B*_*T*) of CG ESCRT-II complex (**Fig. S2**) closely matches the binding affinity of the AA ESCRT-II complex.

### ESCRT-I Oligomerization and ESCRT-I/II Co-Assembly Dynamics

Our prior CG MD simulations demonstrated that higher-order ESCRT-I complexes preferentially oligomerize when the diameter of the inner aperture of the Gag lattice is comparable to the outer diameter (∼50 nm) of the ESCRT-I ring formed by the outward projecting UEV domains (41). This scenario is most likely achieved in the late stages of immature lattice growth when the underlying membrane is significantly deformed (70,71). Here, we extended our previous CG MD simulations to investigate the dynamics of ESCRT-I oligomerization and ESCRT-I/II co-assembly. The system in our CG MD simulations is created such that it mimics a late-stage budding immature virion. Therefore, the system consists of a hemispherical Gag shell wrapped with membrane emulating a narrowing bud neck (**Fig. 2C**). We added multiple copies of ESCRT-I and ESCRT-II to simulate higher-order assembly of upstream ESCRT complexes. The CG model of full-length ESCRT-I consists of the Gag-binding UEV domain, stalk, linker connecting the UEV and stalk, and finally the headpiece region capping the stalk (**Fig. 2*A***). The linker provides flexibility to the UEV, allowing the UEV to diffuse at the Gag lattice surface. The ESCRT-I CG model is assembly competent when templated by the immature Gag lattice; however, it does not undergo oligomerization in solution (41). The CG ESCRT-II is modeled as a membrane associating protein as described in the previous section. We simulated ESCRT-I oligomerization using well-tempered metadynamics (WTMetaD) to estimate the optimal ESCRT-I oligomer size and investigate the differences between the assembly behavior with and without ESCRT-II (72,73). We prepared triplicate simulations for both cases (with and without ESCRT-II), and the simulations were evolved for 200 × 10^6^ CG MD timesteps (*τ*_CG_). In the WTMetaD CG MD simulations, we biased the sum of the coordination number of the largest ESCRT-I oligomer in the system as a collective variable (CV) to enhance ESCRT-I assembly (74). Then we reweighted the trajectories to calculate the probability distribution of the largest ESCRT-I oligomer size observed in our simulations (75,76). For the CG MD simulations with ESCRT-II, a second CV was used to characterize the likelihood of ESCRT-II diffusing to the bud neck. Additional details on the CVs and WTMetaD CG MD simulations are provided in the *Methods* section.

The time profiles of the largest ESCRT-I oligomer in the WTMetaD simulations with and without ESCRT-II are shown in **Fig. 3*A***. The largest ESCRT-I oligomer spontaneously grows and partially disassembles multiple times throughout the course of the simulations, demonstrating the effectiveness of the CV in characterizing the process. The cumulative probability distribution of the largest ESCRT-I oligomer calculated from all the replicate simulations is shown in **Fig. 3*B***. The largest ESCRT-I oligomer typically fluctuates between a 7-mer and 9-mer in simulations without ESCRT-II, constituting ∼ 80% of the largest ESCRT-I oligomers observed in these simulations. The results agree with our previous unbiased CG MD simulations, where we observed growth of up to a 9-mer approaching the 12-mer ring formed by the ESCRT-I headpiece in the crystal structure (41). Furthermore, ESCRT-I oligomer distribution in our simulations lies in the range of 3-11 copies of TSG101 (a subunit of ESCRT-I) imaged in budded HIV-1 particles (77).

**Fig. 3.**
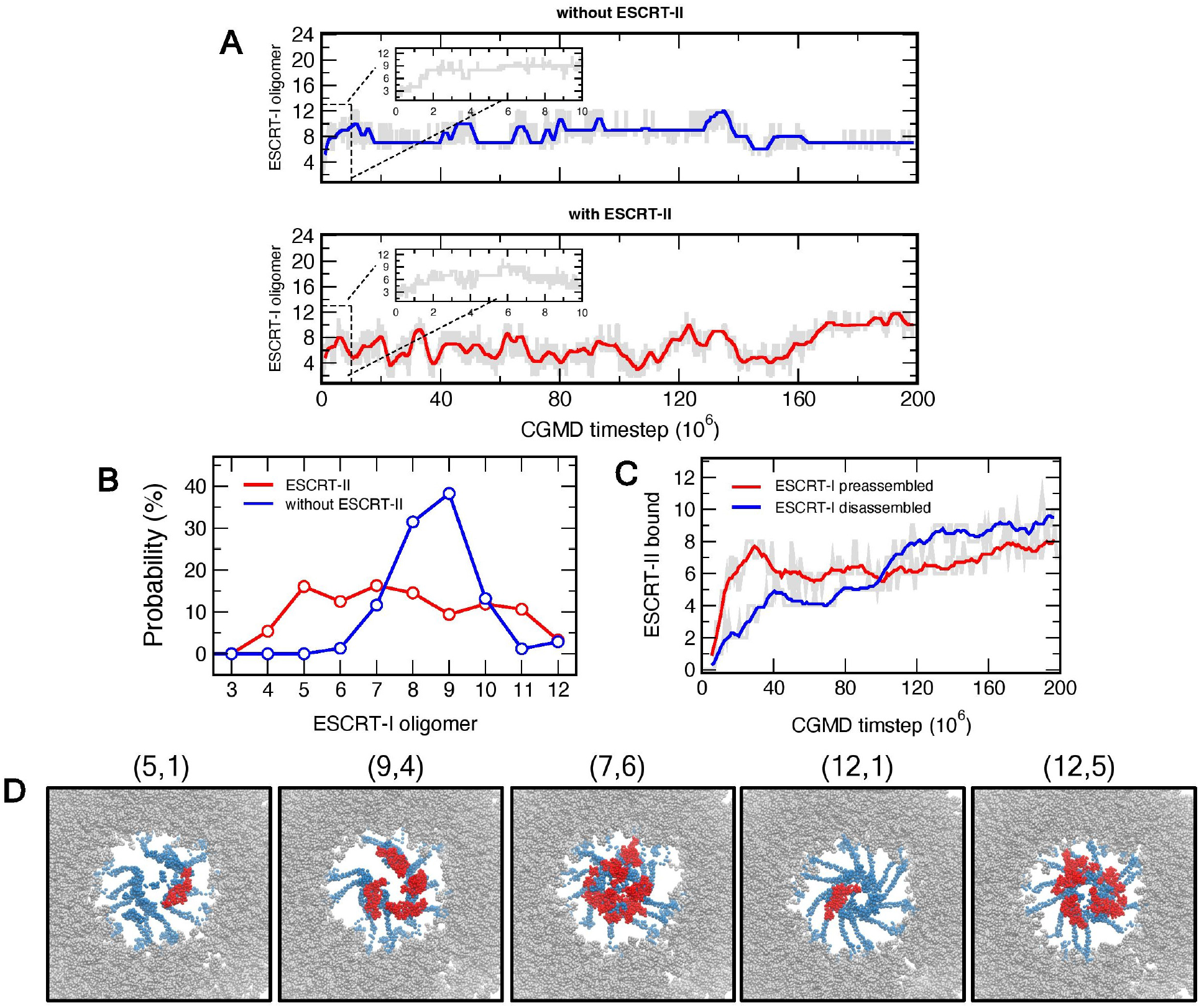
Dynamics of ESCRT-I oligomerization and ESCRT-II recruitment. (*A*) Time series profile of ESCRT-I oligomerization without ESCRT-II (top panel) and with ESCRT-II (bottom panel) from the WTMetaD CG MD simulations. The faded region in gray is the raw data depicting the largest ESCRT-I oligomer. The solid line is the blocked moving average calculated over 1 × 10^6^ timesteps for 200 × 10^6^ timestep long trajectories. (*B*) Probability distribution of the largest ESCRT-I oligomer size in the simulation without ESCRT-II (blue) and with ESCRT-II (red) was presented after reweighting the WTMetaD trajectories. The probability distribution is calculated from final 100 × 10^6^ timesteps of three replicas for each system. (*C*) Time series profile of ESCRT-II recruitment in simulations with ESCRT-I fully disassembled in the initial configuration (solid blue line), and ESCRT-I preassembled to the 12-mer ring in the initial configuration (solid red line). (*D*) Snapshots (top-view looking into the virion from the bud neck) from the simulations of ESCRT-I oligomerization and ESCRT-II recruitment. Gag, ESCRT-I and ESCRT-II are shown in silver, blue and red spheres, respectively. Each snapshot is titled as (*N*_1_, *N*_2_), where *N*_1_ is the largest ESCRT-I oligomer in the system, and *N*_2_ is the number of ESCRT-II copies bound to ESCRT-I. The CG membrane beads are not shown for clarity. The first three snapshots (left to right) correspond to simulations in which the ESCRT-I is fully disassembled in the initial configuration. The final two snapshots correspond to simulations in which the ESCRT-I is preassembled in the initial configuration.

We now discuss the ESCRT-I oligomerization behavior again but in presence of ESCRT-II. The time profiles of ESCRT-II binding to the ESCRT-I oligomers show that 7-9 copies of ESCRT-I/II supercomplexes are formed at the bud neck (**Fig. 3*C* and 3*D***). Our simulations indicate that ESCRT-I is still assembly-competent in the presence of ESCRT-II, with the largest ESCRT-I oligomer fluctuating between 4-mer and 10-mer. In presence of ESCRT-II, however, smaller ESCRT-I oligomers (4-mer to 6-mer) appear to be more prevalent when compared to the simulations of ESCRT-I oligomerization without ESCRT-II. We therefore conjecture that crowding of the bud neck due to localization of multiple ESCRT-II copies may slightly interfere with the ESCRT-I oligomerization dynamics.

To assess the conformational features of the ESCRT-I/II supercomplexes formed at the bud neck we also performed CG MD simulations of ESCRT-II binding to the fully assembled 12-mer ESCRT-I ring. We found that 5-8 copies of ESCRT-II are recruited to the composite headpiece of the ESCRT-I ring which is, importantly, comparable to the results for the simulations starting with fully disassembled ESCRT-I as described in the previous paragraphs. Taken together, the number of ESCRT-II copies bound to ESCRT-I in our simulations is in agreement with a previous model that estimated 6-10 copies of ESCRT-II can be docked in a bud neck of comparable dimension as in the present study (78).

A visual inspection of the ESCRT-I/II supercomplexes formed in the CG MD trajectories reveal extended curved complexes that scaffold the bud neck (**Fig. 4*A*, 4*B***). To characterize the conformation (extendedness or compactness) of the supercomplexes, we calculated the distribution of an end-to-end distance-based metric (*D*_E1-E2_). Here, *D*_E1-E2_ is defined as the distance between the centroid of the ESCRT-I UEV and the C-terminal WH2 domain of ESCRT-II VPS25 subunits. Alternatively, for a particular ESCRT-I/II supercomplex, *D*_E1-E2_ also signifies the spatial distribution of the VPS25 subunits at the bud neck. The peak of the *D*_E1-E2_ distribution spans between 25 and 29 nm, with *D*_E1-E2_ values ranging up to 35 nm (**Fig. 4*B***). In these extended conformations, the bound orientation of the ESCRT-II extends between the ESCRT-I headpiece (or the inner edge of the bud neck proximal to Gag) and outer edge of the bud neck (distal to Gag), the C-terminal tip of the VPS25 positioned closer to the outer edge of the bud neck. Conversely, a minor subpopulation of ESCRT-I/II supercomplexes corresponding to the low *D*_E1-E2_ values represent compact conformations in which the ESCRT-II is pointing downwards towards the Gag lumen.

**Fig. 4.**
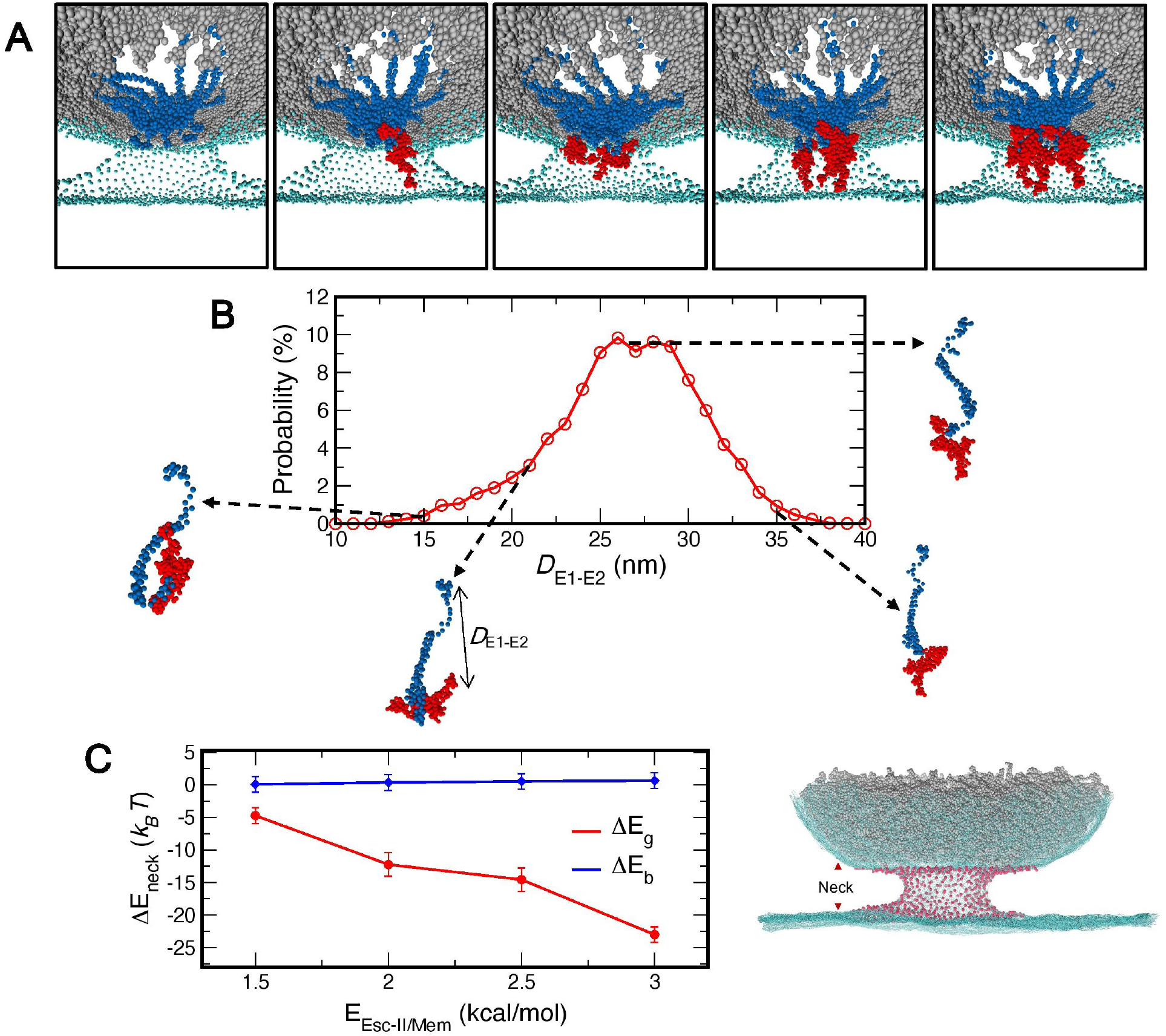
Conformation of the ESCRT-I/II supercomplex. (*A*) Representative snapshots showing ESCRT-II localization at the bud neck and stepwise recruitment of ESCRT-II by the ESCRT-I ring at the bud neck. The leftmost panel depicts the bud neck before ESCRT-II recruitment. The next four panel depicts an ESCRT-II binding event (up to four ESCRT-II copies). The central CG bead of the 3-site CG lipid is shown in cyan spheres. Note that the membrane is sliced vertically in the snapshots to show the interior of the bud neck and the Gag lumen. Gag, ESCRT-I and ESCRT-II are shown in silver, blue and red spheres, respectively. The depicted process is also shown in Supporting movie S1. (*B*) Probability distribution of *D*_E1-E2_ calculated over final 100 × 10^6^ timesteps of each replica simulation. Snapshots show 1:1 ESCRT-I/II supercomplexes. (*C*) The change in membrane bending energy (Δ*E*_neck_) of the bud neck (in our system the bud neck contains ∼550 CG lipids) associated with the formation of the upstream ESCRT complexes plotted as a function of the protein-membrane interaction (*E*_Esc-II/Mem_). The contribution of the bending energy from mean curvature (Δ*E*_b_) and Gaussian curvature (Δ*E*_g_) is shown blue and red, respectively. The right panel shows the CG lipids in bud neck (red spheres). The CG lipids not part of the bud neck is shown in cyan mesh.

### ESCRT-I/II Supercomplex and Bud Neck Curvature

The locus of ESCRT-I binding to ESCRT-II and the membrane targeting sites of ESCRT-II are in close vicinity. When bound to ESCRT-I, the membrane targeting sites of ESCRT-II are distributed contiguous to the ESCRT-I headpiece lining the inner edge of the bud neck (**Fig. S3**). Therefore, binding of multiple copies of ESCRT-II to the headpiece of ESCRT-I ring pins the inner edge of the bud neck to the Gag-ESCRT-I complex, specifically pulling the inner edge of the bud neck closer to the inner ring of the ESCRT-I headpiece. Protein-membrane interactions therefore drive spontaneous change in membrane curvature. It is well known that proteins can induce membrane curvature through scaffolding of the membrane by protein complexes (79-82), membrane association and insertion of hydrophobic helices (83-85), and protein crowding on membrane surfaces (64,86). Based on the higher-order upstream ESCRT assemblies formed at the bud neck we examined how these complexes modulate the curvature of the bud neck. We estimated the change in membrane bending energy (Δ*E*_neck_) of the bud neck associated with the formation of the upstream ESCRT complexes from direct calculation of mean and Gaussian curvatures of the bud neck. **Figure 4*C*** shows that there is a significant decrease in the membrane bending energy of the bud neck in presence of the upstream ESCRT complexes relative to that of free bud neck. The overall change in the membrane bending energy arises predominantly from the decline of the Gaussian bending term (Δ*E*_g_). We note that the contribution (∼25 *k*_B_T) of the upstream ESCRT complexes to the change in bud neck energy is relatively modest when compared to overall change in membrane energy (between 150 to 200 *k*_B_T) required for significant constriction of the bud neck to be comparable to dimensions likely observed in budding HIV-1 virions (66,87-89).

To assess whether the protein-membrane interactions impact the bud neck curvature, we modified the interaction strength (*E*_Esc-II/Mem_) between the lipid targeting sites of ESCRT-II and headgroup of the CG lipid. As the protein-membrane interactions are weakened the change in the Gaussian bending energy due to the assembly of upstream ESCRT complexes gradually diminishes. In other words, for weaker protein-membrane interactions multiple copies of ESCRT-II localize at the bud neck and bind to ESCRT-I without significantly altering the curvature of the bud neck. Our results demonstrate that membrane coating by upstream ESCRT assemblies at the viral budding site can impose negative Gaussian curvature and constrict the bud neck. These results are in agreement with recent experimental observations from electron microscopy images that show that localization of upstream ESCRTs at high density can facilitate membrane deformation (63).

## Discussion

To shed light on the organization of the upstream ESCRT machinery at the HIV-1 budding site we have developed and applied “bottom-up” CG models of Gag, ESCRT-I, and ESCRT-II to study the dynamical mechanisms involved in the formation of the upstream ESCRT assemblies templated by the immature Gag lattice in the emerging virion bud. The CG models were developed from extensive all-atom MD simulations. The CG MD results recapitulate key experimental observations, such as the higher-order oligomerization behavior of ESCRT-I and membrane-targeting behavior of the full-length ESCRT-II complex (41,42). Using these CG models, we have thus investigated the ESCRT-I and ESCRT-II co-assembly and ESCRT-I/II supercomplex formation.

Our CG MD simulations demonstrate key features of the ESCRT-I/II supercomplexes formed at the bud neck. These supercomplexes are typically extended, spanning between the upper edge of the immature Gag lattice to the upper tip of the bud neck. The extended conformations observed in our simulation are in qualitative agreement with conformation of ESCRT-I/II supercomplexes reported in solution (78). The complexes in the extended form are curved, scaffolding the bud neck. The shape of the complexes observed in our simulations is also similar to the columnar assemblies visualized by Electron Microscopy at the site of Gag immature (56,71). Furthermore, these complexes also resemble other membrane scaffolding proteins such as the crescent-shaped Bin/Amphiphysin/Rvs (BAR) domain proteins (90,91), and *α*-synuclein (92).

We note that in the late stages of the HIV-1 Gag assembly when immature Gag lattice has grown past a hemisphere, the bud neck can be approximated as catenoid-shaped (93). Catenoid-shaped bud necks are characterized by negative Gaussian curvature, and near-zero mean curvature. We find that the neck coating upstream ESCRT assemblies scaffolded by the immature Gag lattice further reinforce negative Gaussian curvature, therefore contributing to neck constriction. Our analysis of the membrane bending energy of the bud neck in our simulations demonstrates that the upstream ESCRT assemblies contribute to the overall change in membrane bending energy required for bud neck constriction estimated from scaling theories of neck constriction (89). The conformation of the ESCRT-I/II supercomplexes also have important implications for ESCRT-III nucleation. In the extended conformation, the VPS25 subunits are distributed in the vicinity of the upper edge of the bud neck and expected to be accessible for CHMP6 binding. These ESCRT-II:(CHMP6)_2_ complexes can then act as nucleation seeds (46), with the ESCRT-III filaments nucleating and propagating outwards from the upper edge of the bud neck to the flat section of the membrane surrounding the viral budding site.

In this study we considered the organization of the upstream ESCRT-I and ESCRT-II complexes. The upstream ESCRT machinery also consists of ALIX which offers an alternate pathway for ESCRT-III recruitment. ALIX consists of the N-terminal BRO1 domain (residues 1-359), central V domain (360-716) and the C-terminal proline-rich domain (717–868). Gag recruits ALIX through interactions between the YPX_n_L motifs in the late domain of the Gag and second arm of the V domain (40,94) and between the BRO1 domain and nucleocapsid (NC) domain of the Gag mediated by RNA (95). It remains an open question whether ALIX monomer adopts a “closed” or “open” conformation to simultaneously bind to both the YPX_n_L motif in the p6 domain and NC domain of Gag (96). We anticipate that a follow-up atomistic simulation study combining a Gag hexamer bundle, RNA, and ALIX can be designed to determine the binding mode and preferred conformational state of ALIX. Based on the results of these atomistic simulations we can derive a bottom-up CG model of ALIX to incorporate interactions between ALIX, Gag, and RNA. These bottom-up CG models can then be used to investigate ALIX recruitment and organization by the immature Gag lattice mediated by RNA at the budding site.

Finally, the “budding virion” CG model presented in this work consists of the Gag, upstream ESCRT-I, ESCRT-II, and lipid. The model will be continuously updated to add new components such as ALIX and RNA, and the interactions between all the components of the virion will be refined as new structural and biochemical data becomes available. Our future research efforts will also focus on developing and applying a “bottom-up” CG model of ESCRT-III to investigate the dynamics of upstream ESCRT-mediated nucleation and growth of ESCRT-III polymers and bud neck constriction driven by these polymers. A key outstanding issue that remains to be explored is whether ESCRT-I/II or ALIX branch of the upstream ESCRT machinery is the dominant pathway or if both branches contribute in a concerted manner to the recruitment and nucleation of ESCRT-III polymers during HIV-1 budding. Our updated “budding virion” CG model will strive to answer these questions.

## Methods

### Atomic-level protein models

The initial atomic model for the full-length ESCRT-II complex was constructed by combining the crystallographic fragments of the VPS22:VPS36 complex (PDB: 3CUQ, 2ZME), VPS25 (PDB: 3CUQ, 2ZME) and GLUE domain (PDB: 2HTH) (42,97). The missing N-terminal *α*-helix (VPS22: 1-33) and the linker (VPS36: 132-171) connecting the C-terminal region of the GLUE domain and VPS36 core were constructed using the amino acid sequence from the Robetta server (98,99).

### All-Atom (AA) MD simulations

The membrane model used in this study to investigate the conformational state of membrane-bound ESCRT-II was composed of 76% POPC, 20% POPE, and 4% PIP2, based on the experimental study of ESCRT-II membrane binding (42). Two symmetric membrane models of the above-mentioned composition of dimension 17.2 × 17.2 nm^2^ and 9.5 × 9.5 nm^2^ were built to study membrane binding of ESCRT-II complex and monomeric membrane targeting domains, respectively. The smaller lipid bilayer system was used to perform umbrella sampling (US) enhanced free energy simulations. The model membranes were created and solvated using CHARMM-GUI Membrane Builder (100). The solvated bilayers were minimized, relaxed using the standard six-step protocol provided on CHARMM-GUI for a cumulative 750 ps, and then equilibrated for additional 200 ns. The coordinates of the lipid molecules were then extracted from the final configuration of the equilibrated trajectories and merged with the protein coordinates. The ESCRT-II complex was oriented such that in the initial configuration the highly basic VPS22-H0 helix was 1 nm away from the membrane surface and the H0-helix was coplanar to the membrane surface. The composite membrane-protein system was then equilibrated by applying harmonic positional restraints (spring constant value of 239 kcal/mol/nm^2^) on protein and lipid heavy atoms for 500 ps, followed by another 500 ps of equilibration by harmonic restraints on only the protein C_a_ atoms. A further 300 ps of restrained equilibration was performed, and configurations were saved every 100 ps to be used as initial structures for independent production runs. The restrained simulations were performed with a constant NVT ensemble. The temperature of the system was maintained at 310 K using stochastic velocity rescaling thermostat with a time constant of 1 ps. Production runs were carried out for 2000 ns in constant NPT ensemble at 310 K and 1 bar. The temperature was maintained using Nose-Hoover chain thermostat with a 2 ps time constant, and pressure with semi-isotropic (*x* and *y* directions coupled) Parrinello-Rahman barostat with a 10 ps time constant. Identical setup protocol was used to build and equilibrate the protein-membrane system of the monomeric GLUE domain (VPS36: 1-132) and VPS22_Mem_ (VPS22: 1-76) for the US simulations, and VPS28-CTD docked to VPS22:VPS36 subcomplex to derive the ESCRT-I/II CG attractive interactions.

All simulations were performed with periodic boundary conditions in *x, y* and *z* directions, and timestep of 2 fs. The protein and lipid were modeled with CHARMM36m force field (101), and water was modeled with TIP3P parameters (102). LINCS algorithm was used to constrain the bonds between heavy and hydrogen atoms (103). Electrostatic interactions were computed using the particle mesh Ewald method (104), and van der Waals force was truncated smoothly to zero between 1.0 and 1.2 nm. All simulations were performed using Gromacs 2019 package (105).

### AA MD Umbrella Sampling Simulations

We performed umbrella sampling simulations to estimate the free energy of binding to the membrane for the monomeric GLUE domain and VPS22_Mem_. The distance between the *z* component of the center of mass of membrane bilayer and protein was used as the reaction coordinate for the US simulations. To generate configurations for US simulation windows, we performed steered molecular dynamics (SMD) simulations for each system. The initial configuration for the SMD simulations corresponds to the equilibrated structure at the end of 200 ns production run for each system. In each case, the protein was pulled from the equilibrated bound state to the unbound state (6 nm from the membrane center). SMD simulations were performed with a constant pulling velocity of 10^−4^ nm ps^-1^ and configurations were saved every 0.05 nm to be used as starting configuration for umbrella sampling windows. We note that in the SMD simulations 1 fs timestep is used. In the SMD simulations, harmonic positional restraints (spring constant value of 239 kcal/mol/nm^2^) were applied to the lipid heavy atoms to prevent upward lipid displacement. For the US simulations, constrained simulation at each umbrella window was performed with 1000 kcal/mol/nm^2^ harmonic force constant. For each protein, total 42 umbrella sampling windows were used to compute the PMF profile. Each umbrella window was evolved for 100 ns, resulting in a cumulative simulation time of 8400 ns. Potential of mean force (PMF) profile was computed from the last 75 ns of each umbrella sampling window using the weighted histogram analysis method (WHAM) (106,107). The statistical uncertainty of the PMF was evaluated using the block averaging method by dividing the data into five blocks of 15 ns each, and PMF was calculated for each block (108). The standard deviation of the PMF was calculated from the five PMF’s corresponding to each block. The histogram showing the overlap between consecutive umbrella windows is shown in **Fig. S4**.

### CG model generation

We derived a bottom-up model of ESCRT-II from the AA MD trajectories of the ESCRT-II bound to the membrane. The CG sites are mapped from the atomistic trajectories using Essential Dynamics Coarse-Graining (EDCG) method (109). In the EDCG method, contiguous C_*α*_ atoms of the protein were grouped to CG sites to reflect the atomistic collective fluctuations, specifically the motions in the essential subspace determined from principal component analysis of the reference all-atom trajectories. The resulting CG model of the ESCRT-II has 222 sites: the VPS22 and VPS36 subunit have 54 and 94 CG sites respectively, and the two VPS25 subunits have 37 sites each. After defining the CG mapping, intramolecular interactions between the CG sites were represented as a network of effective harmonic bonds between a central CG site and all other CG sites within 3 nm. The force constants of the harmonic bonds are derived using the hetero-elastic network model (hENM) method (110). The final 500 ns of the all-atom trajectory (replica1) was used to define the CG mapping and harmonic spring constants. Intermolecular CG interactions between proteins were modeled using a combination of excluded volume (*E*_excl_) to avoid unphysical overlap between CG beads and attractive interactions (*E*_attr_). For *E*_excl_, a soft cosine potential, 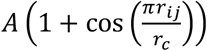 is used, *r*_*ij*_ is the pairwise distance between CG site types *i* and *j*. The value of *A* is 25 kcal/mol for all *ij* pairs. The distance cutoff (*r*_c_) between excluded volume interactions was set at 2.5 nm. Attractive interactions (*E*_attr_) were modeled as a pairwise Gaussian potential, 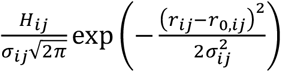. Here *r*_*0,ij*_ and *σ*_*ij*_ are the mean and standard deviation of the distance between CG site types *i* and *j* determined by fitting to the corresponding pair correlation through least-squares regression. All pair potentials used a 2.5 nm radial cutoff, except for the attractive interactions to drive ESCRT-I oligomerization where a 3 nm radial cutoff was used.

The attractive pairwise interaction between VPS28-CTD of ESCRT-I and ESCRT-II was derived from an AA MD simulation trajectory of VPS28-CTD complexed to a membrane bound VPS22:VPS36 subcomplex. The initial configuration of this simulation was prepared by docking VPS28-CTD to the VPS36 linker (42,43). The last 500 ns of this trajectory was used to map the C_*α*_ atoms to corresponding CG sites. For the VPS22:VPS36 subcomplex identical mapping is used as in the full-length ESCRT-II complex described previously. For the VPS28-CTD CG sites are mapped using the EDCG method with an average resolution of ∼8 amino acid residues per CG site (109). The CG mapping resolution is consistent with the CG model of ESCRT-I headpiece, UEV domain and stalk as in our prior CG study (41). From the CG mapped trajectory of VPS28-CTD complexed to a membrane bound VPS22:VPS36 subcomplex, we identified CG sites of VPS28-CTD (*i*) within 2 nm of any CG sites (*j*) of the VPS22-VPS36 subcomplex, and with a standard deviation lower than 0.18 nm. These CG site pairs (*ij*) are assumed to be in close contact and hence contributing towards ESCRT-I/II association. The coefficient *H*_*ij*_ of these CG pair sites were optimized through the relative-entropy minimization (REM) method (111). A detailed description of the REM method to derive the CG attractive interprotein interactions from AA MD trajectories is provided in ref. (112,113), where REM was used to optimize attractive interactions between SARS-CoV-2 structural proteins.

To model the ESCRT-I headpiece, UEV, and stalk we used the same CG mapping and interaction parameters as in our previous study of ESCRT-I oligomerization templated by immature Gag lattice (41). To allow ESCRT-I/II association the first particle of CG VPS28-CTD was attached to the last particle of VPS28-NTD in the headpiece through a harmonic bond of force constant 50 kcal/mol/nm^2^ and equilibrium distance 0.15 nm. The updated ESCRT-I CG model contains 109 CG sites. The attractive interactions driving ESCRT-I oligomerization is modeled with pairwise Gaussian potential. The interaction parameters are described in ref. (41). Briefly, the value of *H*_*ij*_ (−0.9 nm kcal/mol) is chosen such that the ESCRT-I molecules do not oligomerize in the solution. However, the value of *H*_*ij*_ is strong enough to allow extensive oligomerization when templated by the immature Gag lattice of appropriate diameter. The CG model of the Gag polyprotein consists of matrix (MA), capsid/spacer peptide 1 (CA/SP1), and nucleocapsid (NC) domains. To model the association of the ESCRT-I UEV domain and p6 domain of Gag, a CG binding site was added to the C-terminal end of the NC domain. Attractive interactions are added between the Gag binding site and final two C-terminal CG sites of UEV domain and the Gag binding site. The details of the Gag CG model generation and ESCRT-I/Gag interactions are described in ref. (41).

All membrane association interactions between the protein CG sites (Gag and ESCRT-II) and CG lipid head group were modeled using a 12-6 Lennard-Jones potential (*E*_sclj_) with a modified soft-core (114), 4*ελ*^*n*^ [1⁄(*α*_*LJ*_(1 − *λ*)^2^ + (*r*⁄*σ*)^6^)^2^ − 1⁄(*α*_*LJ*_ (1 − *λ*)^2^ + (*r*⁄*σ*)^6^)]. Here, *n* = 2, *α*_*LJ*_ = 0.5, *λ* = 0.6 and *σ* = 1.5 nm. Gag interacts with the lipid head group through three beads in the N-terminal region of the MA domain with an interaction strength, *ε* = 2.5 kcal/mol. ESCRT-II interacts with the lipid head group with an optimal interaction strength, *ε* = 3.0 kcal/mol. We note that in the main text, membrane association strength of ESCRT-II (*E*_Esc-II/Mem_) is identical to *ε*. For ESCRT-II, we also explored lower interaction strengths varying *ε* (*E*_Esc-II/Mem_) from 1.5 kcal/mol to 2.5 kcal/mol.

The CG atom type index of the composite system is listed in Table S1. The details of the attractive interactions that drive ESCRT-I/II association and ESCRT-I oligomerization are listed in Table S2 and S3 respectively. Additional details on the CG model bonding topology, hENM force constant, equilibrium bonding distances and the coordinates of the initial coarse-grained models are listed in https://github.com/arpahudait/Gag-ESCRT_budding.

### CG MD System Setup and Simulation Settings

We prepared the ‘*bud neck’* model system resembling the budding site of the immature HIV-1 virion with the same protocol as our previous CG study (41). Briefly, we prepared model Gag shells by extracting coordinates of the Gag polyprotein assemblage from cryo-electron microscopy maps of immature HIV-1 lattices. The diameter (*D*_Gag_) of the inner aperture of the Gag shell for all systems in this study is 50 nm. This is the optimal dimension for the 12-membered ESCRT-I ring formation (41). The extracted Gag lattice was then mapped to CG Gag and wrapped with a membrane surface that resembles a budding neck. The final lateral dimension of the simulation cell was 136 nm × 136 nm in the *x, y* directions respectively. The system was then equilibrated for 10 × 10^6^ CG timesteps by integrating only the CG lipids (the protein is held fixed) to allow relaxation of the lipid sheet, neck, and lipid around Gag. The 12-membered ESCRT-I ring was then manually docked to the Gag shell. The system was then further equilibrated for 20 × 10^6^ CG timesteps. In these simulations, the CA/SP1 domain of the Gag is held fixed, while other domains of the Gag, ESCRT-I and lipids are integrated. We found that in 20 × 10^6^ CG timesteps, the UEV domain of all 12 ESCRT-I molecules binds to the Gag. To generate initial configuration for the simulations of ESCRT-I oligomerization, the attractive interactions between the ESCRT-I proteins that drive oligomerization were turned off and the system is evolved for 50 × 10^6^ CG timesteps. During this simulation, the ESCRT-I ring completely disassembles and the ESCRT-I molecules randomly diffuses in the Gag lumen generating randomized initial configurations. To summarize, we generated two systems – 1. ESCRT-I is fully assembled to a 12-membered ring, and 2. 12 ESCRT-I monomers are bound to the Gag lattice, however fully disassembled. To prepare simulations with ESCRT-II, 16 ESCRT-II proteins were initially arranged in a two-dimensional uniform grid and placed in the *xy* plane 2 nm above the membrane surface. Additionally, the initial ESCRT-II grid was prepared such that the distance between the Gag shell center and ESCRT-II proteins was less than 60 nm. Finally, the system was equilibrated for 20 × 10^6^ CG timesteps by turning off the ESCRT-I/II interactions to allow the ESCRT-II proteins to bind to the membrane.

CG MD simulations were performed using the LAMMPS MD software interfaced with PLUMED (version 2.7) (115,116). In all CG MD simulations, the equations of motion were integrated with the Velocity Verlet algorithm using a time step (*τ*_CG_) of 50 fs. Periodic boundary conditions were used in the *xy* directions, while in the *z* direction the simulation cell was non-periodic. The simulations were performed in the constant *Np*_*xy*_*T* ensemble. The pressure in the *xy* direction was controlled at 0 bar with the Nose-Hoover barostat using a coupling constant of 2000*τ*_CG_ (117). The temperature of the simulation was maintained at 300 K using a Langevin thermostat using a coupling constant of 1000*τ*_CG_ (118). Simulation trajectory coordinates were saved every 1 × 10^6^ CG MD timesteps.

### CG Metadynamics Simulations

We performed well-tempered metadynamics simulations to facilitate oligomerization of ESCRT-I and binding of ESCRT-II to ESCRT-I at the bud neck. We performed two sets of metadynamics simulations: (*i*) Investigating ESCRT-I oligomerization dynamics (with and without ESCRT-II), and (*ii*) ESCRT-II binding to the fully oligomerized 12-membered ESCRT-I ring. To facilitate oligomerization of ESCRT-I we used the collective variable (CV1) reported in ref (74). To define CV1, we first represent each ESCRT-I monomer by the geometric center of the CG sites corresponding to headpiece region. This choice is justified since ESCRT-I oligomerization is driven by interactions at the headpiece region (41). To assess the overall connectivity between the ESCRT-I oligomers, we first calculated the coordination number and then a symmetric adjacency matrix using the coordination number. Both coordination number and elements of the adjacency matrix were calculated using a cubic harmonics function with a radial cutoff of 3.5 nm, while the function decays from one to zero between 3.5 nm and 4 nm (119,120). Then, oligomer sizes were determined by applying the depth first search (DFS) clustering algorithm to the adjacency matrix. Finally, the value of CV1 is the sum of the coordination numbers for all the ESCRT-I molecules in the largest cluster determined by the DFS clustering protocol. To define CV2, we first represented each ESCRT-I and ESCRT-II by the geometric center of the CG sites corresponding to VPS28-CTD and VPS36 linker respectively. Then, the distance (*D*_ij_) between all ESCRT-I/II pairs were calculated. Finally, CV2 is defined as the maximum of all *D*_ij_ values. For simulations of ESCRT-I oligomerization in the presence of ESCRT-II the biases were deposited along CV1 and CV2. For simulations of ESCRT-II binding to the 12-membered ring biases were deposited along CV2. To improve computational efficiency a weak harmonic restraint of force constant 2390 kcal/mol/nm^2^ was applied to prevent CV2 exceeding 60 nm. In the CG metadynamics simulations, the added Gaussian biases had a width of 0.1 (CV1) and 0.02 nm (CV2). The height of the Gaussian biases was set to 0.15 kcal/mol and was deposited every 500 *τ*_CG_ with a bias factor of 100.

### CG Umbrella Sampling Simulations

To assess the binding free energy between ESCRT-II and membrane, we used umbrella sampling (US) simulations. US simulations were performed as a function of the distance between the center of membrane and ESCRT-II. To calculate the center of mass of ESCRT-II only the CG sites of VPS22:VPS36 subcomplex was considered, since the VPS25 subunits do not associate with the membrane. We restrained the distance at each umbrella window with a harmonic force constant of 200 kcal/mol/nm^2^. In the US simulations, the dimension of the membrane surface was 50 nm × 50 nm in the *x, y* directions. The distance between the membrane center and ESCRT-II was varied from 3.2 to 18.2 nm generating 150 windows at 0.1 nm. For each window, simulations were evolved for 2 × 10^6^ CG MD timesteps. The potential of mean force (PMF) profile was computed from the final 1.5 × 10^6^ CG MD timesteps of each umbrella sampling window using the weighted histogram analysis method (WHAM) (106,107). US simulations were performed by varying *E*_Esc-II/Mem_ from 1.5 kcal/mol to 3.0 kcal/mol at intervals of 0.5 kcal/mol. For each *E*_Esc-II/Mem_ value, three independent US simulations were generated. The error bar for the binding affinity was calculated from the three PMF profiles at the umbrella window corresponding to the minima.

### Analysis

To identify protein-protein association we used the following distance-based criteria:

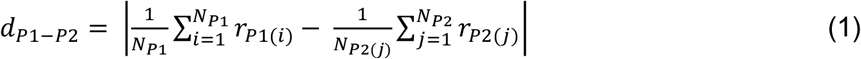

where *P*1(*i*) and *P*2(*j*) are the indices of the CG site types used for the calculation in protein P1 and P2, respectively. *r*_P1(*i*)_ and *r*_P2(*j*)_ denote coordinates of the CG sites used for the calculation. *N*_P1_ and *N*_P2_ are the number of CG sites used for the calculation in protein *P*1 and *P*2. *d*_P1-P2_ is the distance between a specific protein pair. To determine ESCRT-I/II association, we considered CG site types 87-90, 94, 98-99 of ESCRT-I and CG site types 81,89-92 of ESCRT-II. A protein pair was classified as associated if *d*_P1-P2_ is less than 3.5 nm.

To calculate membrane curvature and bending energy, we first constructed a surface mesh from the position of the central CG lipid by using the Delaunay triangulation method (121). In the triangulated surface mesh, each vertex (*V*) forms multiple (*T*) triangles with the neighboring vertices. The vertex area (*A*_i_) was then defined as 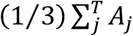, where *A*_j_ is the area of each triangle, and the sum is over the all the *T* triangles that the vertex *V* comprises. To create the triangulated mesh, all lipid molecules within 6.5 nm of the central lipid were considered. The mean (*H*) and Gaussian curvature (*K*) was then calculated from the triangulated surface mesh using the protocol reported in ref. (122). The contribution of the mean curvature to bending energy was calculated as 2*κ*_*B*_*A*_*i*_*H*^2^. The contribution of the Gaussian curvature to bending energy is calculated as *κ*_*G*_*A*_*i*_*K*. Here, *κ*_*B*_ and *κ*_*G*_ are the mean and Gaussian bending rigidity modulus, respectively. The value of the mean bending rigidity modulus (*κ*_*B*_) is 12.2 *k*_*B*_T (69). The Gaussian bending rigidity modulus was approximated as 0.9*κ*_*B*_ based on ref. (123).

## Data Availability

The CG models of the Gag, ESCRT-I, ESCRT-II, and CG lipid are available in the Github repository (https://github.com/arpahudait/Gag-ESCRT_budding). Additional details are provided within the article or Supporting Information. Additional data is available upon request to the author.

## Acknowledgements

The computational resources used in this work were provided by the Research Computing Center (RCC) at The University of Chicago and the Texas Advanced Computing Center (TACC) at The University of Texas at Austin. Simulations were performed using resources provided by Extreme Science and Engineering Discovery Environment (XSEDE) (124), supported by the National Science Foundation grant number ACI-1548562, and Frontera (at TACC) funded by the NSF (OAC-1818253).

## Author contributions

A.H., J.H.H., and G.A.V. designed research; A.H. built systems, performed simulations, analyzed data, created figures, and prepared the initial manuscript draft. All authors contributed to the writing and editing of the final manuscript.

## Funding and additional information

This research was supported by the National Institute of Allergy and Infectious Diseases (NIAID) of the National Institutes of Health (NIH) under grant P50-AI150464 for the Center for the Structural Biology of Cellular Host Elements in Egress, Trafficking, and Assembly of HIV (CHEETAH) (A.H. and G.A.V.), the National Institute of General Medical Sciences (NIGMS) (grant R01-GM063796) (A.H. and G.A.V.), and by NIAID R37-AI112442 (J.H.H.).

## Abbreviations

The abbreviations used are:

ESCRT: endosomal sorting complexes required for transport
AA MD: All-Atom Molecular Dynamics
CG MD: Coarse-Grained Molecular Dynamics
RMSF: root mean square fluctuation

