## Supplemental File for "Organization of Upstream ESCRT Machinery at the HIV-1 Budding Site"

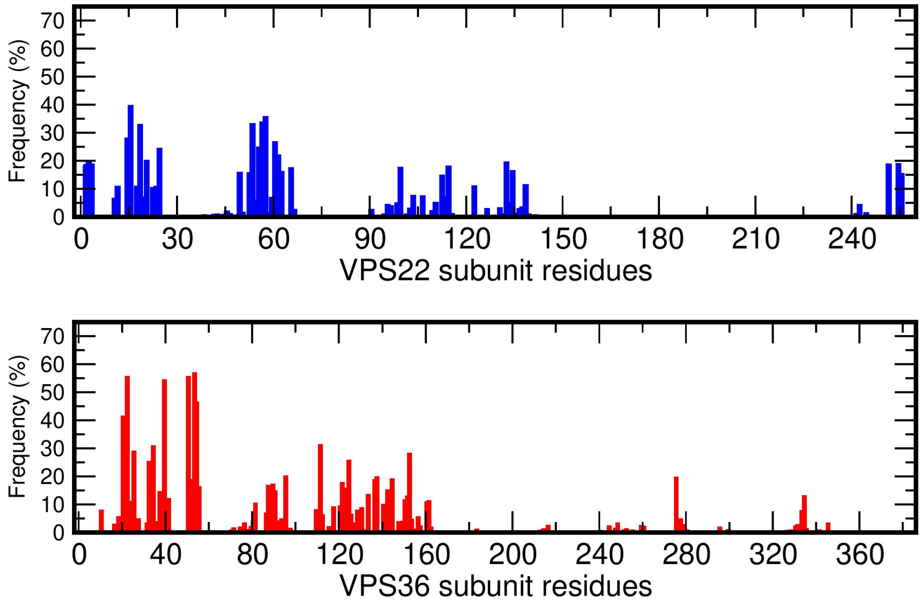


**Fig. S1.** Percentage of MD frames a residue of VPS22 subunit (upper panel), and VPS36 subunit (lower panel) was in contact with PIP2 lipids. The contacts are calculated for residues within 5 Å of PIP2 lipids.

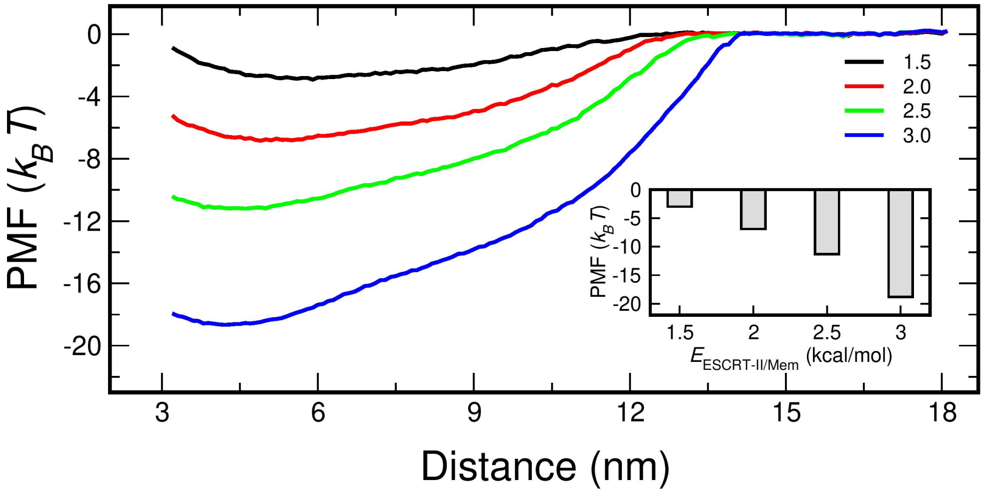


**Fig. S2.** Potential of mean force (PMF) for CG ESCRT-II complex binding to the membrane. From top to bottom each line represents a different $E_{\mathrm{Esc\text{-}II/Mem}}$ (attractive interaction between head group of CG lipid and membrane targeting CG site of ESCRT-II). Inset panel shows the binding affinity (value at the minimum of the curve) for each $E_{\mathrm{Esc\text{-}II/Mem}}$.

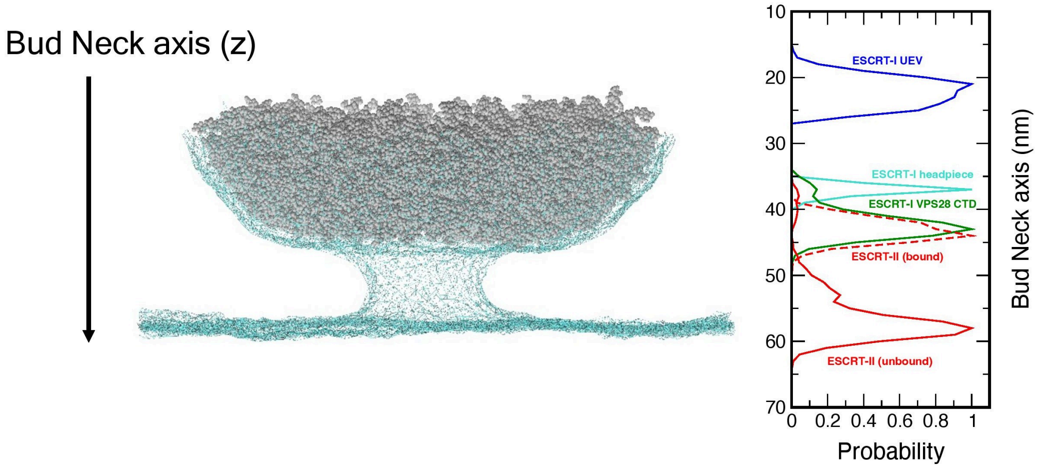


**Fig. S3.** Probability distribution of the different components of the system along the bud neck axis. In our simulations, z axis is the bud neck axis (left panel). Right panel shows the probability distribution along the bud neck for ESCRT-I UEV domain, headpiece region, and VPS28 CTD, and membrane targeting sites of ESCRT-II. In the plot, ESCRT-II (bound) and ESCRT-II (unbound) indicates ESCRT-II proteins that are bound and not bound to the ESCRT-I VPS28 CTD respectively. All probability distributions were calculated from the z coordinates of the selected CG site types. The dimension of the simulation cell is 90 nm in the z direction.

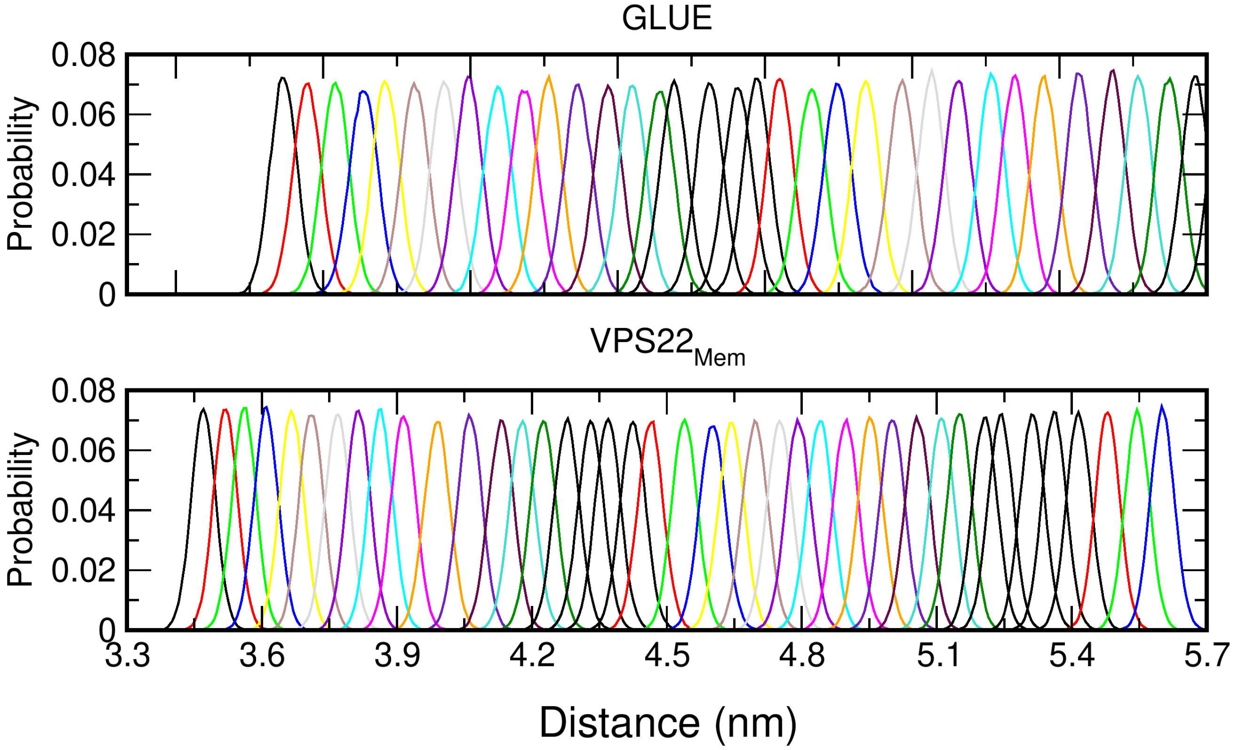


**Fig. S4.** Histogram of the distances between the lipid bilayer center and protein center calculated for consecutive windows in the all-atom Umbrella Sampling simulations for the GLUE and $VPS22_{Mem}$. The histograms are calculated with a bin width of 0.005 nm.

**Table S1**

CG atom types in the composite systems (Gag, CG lipid, ESCRT-I and ESCRT-II):

| **CG type index** | **Protein/Lipid** |
|---|---|
| 1-69 | Gag polyprotein[a,d] |
| 70-72 | 3-site CG lipid |
| 73-171 | ESCRT-I[b] |
| 172-393 | ESCRT-II[c,e] |

a. The matrix (MA), capsid and spacer peptide 1 (CA-SP1), and nucleocapsid (NC) domain consists of CG site type 1-20, 22-56, and 58-68 respectively.
b. TSG101 (UEV), headpiece and VPS28-CTD consists of CG site types 73-91, 126-158, 159-171 respectively.
c. The CG site types corresponding to VPS22, VPS36 and the two VPS25 subunits are 172-225, 226-318 and 319-393.
d. CG site type 4-6 of the MA domain of Gag have attractive membrane-binding interaction with the CG lipid head bead (type 72).
e. CG site type 176-177, 186-187, 229-230, 232-234, 238, 254, 263 of ESCRT-II have membrane-binding interaction with the CG lipid head bead (type 72).

**Table S2**

CG site types with attractive Gaussian interactions to allow ESCRT-I and ESCRT-II association:

| ESCRT-I CG type | ESCRT-II CG type | $H_{ij}$ (nm kcal/mol) | $r_{0,ij}$ (nm) | $\sigma_{ij}$ (nm) |
|---|---|---|---|---|
| 170 | 261 | -0.8 | 0.831 | 0.057 |
| 170 | 262 | -0.8 | 1.059 | 0.063 |
| 160 | 262 | -0.8 | 0.730 | 0.058 |
| 161 | 262 | -0.8 | 1.155 | 0.053 |
| 162 | 262 | -0.8 | 0.975 | 0.137 |
| 160 | 263 | -0.8 | 1.083 | 0.150 |
| 166 | 260 | -0.8 | 0.861 | 0.181 |
| 166 | 261 | -0.8 | 0.923 | 0.138 |
| 171 | 252 | -0.8 | 1.026 | 0.173 |
| 160 | 199 | -0.8 | 0.842 | 0.110 |
| 159 | 201 | -0.8 | 0.866 | 0.096 |
| 160 | 200 | -0.8 | 0.944 | 0.108 |

**Table S3**

CG site types with attractive Gaussian interactions to allow ESCRT-I oligomerization at the headpiece region.

| ESCRT-I CG type | ESCRT-I CG type | $H_{ij}$ (nm kcal/mol) | $r_{0,ij}$ (nm) | $\sigma_{ij}$ (nm) |
|---|---|---|---|---|
| 131 | 134 | -0.9 | 1.800 | 0.09 |
| 131 | 149 | -0.9 | 1.275 | 0.09 |
| 150 | 151 | -0.9 | 1.570 | 0.09 |
| 150 | 152 | -0.9 | 1.744 | 0.09 |
| 152 | 153 | -0.9 | 1.275 | 0.09 |

**Supporting Movie S1**

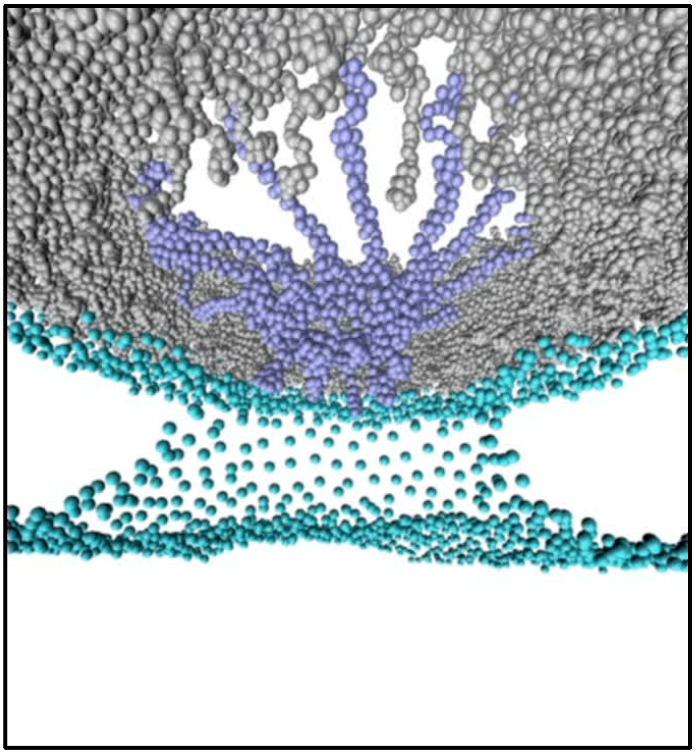
